# Mutualism outcome across plant populations, microbes, and environments in the duckweed *Lemna minor*

**DOI:** 10.1101/448951

**Authors:** Anna M. O’Brien, Jason Laurich, Emma Lash, Megan E Frederickson

## Abstract

The picture emerging from the rapidly growing literature on host-associated microbiota is that host traits and fitness often depend on complex and interactive effects of host genotype, microbial interactions, and abiotic environment. However, testing these main and interactive effects typically requires large, multi-factorial experiments and thus remains challenging in many systems. Furthermore, most studies of plant microbiomes focus on terrestrial hosts and microbes. Aquatic habitats may confer unique properties to plant micriobiomes. We grew different populations of duckweed (*Lemna minor*), a floating aquatic plant of increasing popularity in freshwater phytoremediation, in three microbial treatments (adding no, “home”, or “away” microbes) at two levels of zinc, a common water contaminant in urban areas. Thus, we simultaneously manipulated plant source population, microbial community, and the abiotic environment, and measured both plant and microbial performance as well as plant traits. Although we found little evidence of interactive effects, we found strong main effects of plant source, microbial treatment, and zinc on both duckweed and microbial growth, with significant variation among both duckweed and microbial communities. Despite strong growth alignment between duckweed and microbes, zinc consistently decreased plant growth, but increased microbial growth. Furthermore, as in recent studies of terrestrial plants, microbial interactions altered a duckweed phenotype (frond aggregation). Our results suggest that the duckweed source population, its associated microbiome, and the contaminant environment may all need to be considered in real-world phytoremediation efforts. Lastly, we propose that duckweed microbes offer a robust experimental system for study of host-microbiota interactions under a range of environmental stresses.

## Introduction

Plant performance in harsh environments often depends on the associated microbial community. Plant-microbe interactions commonly alter plant development and trait expression in ways that affect plant fitness, especially under stress (Friesen et al., 2011; Goh et al., 2013). Increasing complexity, interactions between environments and microbes (Smith and Read, 2008; Zhu et al., 2009; Johnson et al., 2010; Lau and Lennon, 2012; O’Brien et al., 2018), plant genotypes and microbes (Johnson et al., 2010; Wagner et al., 2014; Rúa et al., 2016), and three-way interactions among plant genotypes, microbes, and environments (Johnson et al., 2010; Wagner et al., 2014) can all contribute to plant phenotypes and fitness. Indeed, local adaptation and host-microbe coevolution require genotype-by-environment (G x E) and genotype-by-genotype (G x G) interactions for plant or plant *and* microbial fitness, respectively. The geographic mosaic theory of coevolution predicts that three-way interactions (G x G x E) will be widespread (Thompson, 2005), fundamentally shaping plant and microbial evolution. Yet in many systems, testing for G x E, G x G, and G x G x E effects is challenging because of the multi-factorial nature of the necessary study design, the high level of replication required, the relatively long generation times of plants, and the difficulties inherent in measuring microbial “fitness”.

Here, we aimed to develop a novel experimental system to test for interactions among plant genotype, microbial community, and environment using field-collected plants and the microbes with which they associate (and potentially coevolve) in nature. As a focal plant, we chose the tiny, aquatic angiosperm *Lemna minor* (common duckweed) because it reproduces primarily by budding, making it easy to propagate. *L. minor* ocassionally flowers and sets seeds, and it has a higher outcrossing rate and greater genetic diversity than other duckweed species such as *Spirodela polyrrhiza* (Ho, 2017). Thus, *L. minor* collected from different locations are often different clonal genotypes, potentially hosting divergent microbial communities or adapted to different environmental conditions. Furthermore, duckweed very rapidly increase in density under both natural and experimental conditions, facilitating fast experiments. *Lemna minor* is one of the world’s smallest angiosperms, allowing us to grow it in merely 2.5 mL of liquid media and to load 24 experimental replicates on a single standard-size well plate.

Our current understanding of plant-microbe interactions largely comes from studies of nutritional symbioses in terrestrial plants, such as legume-rhizobium or plant-mycorrhizal associations that alleviate a nutrient stress of the plant. However, plants support a diverse community of microbes on their roots (Bulgarelli et al., 2012; Lundberg et al., 2012; Ishizawa et al., 2017b) that can modulate effects of other plant stressors, including environmental contaminants. Microbes can reduce the impacts of contaminants on plants by preventing contaminants from being taken up, by helping plants tolerate or sequester compounds, or by promoting plant growth in general and simply diluting harm done by the contaminant (Rajkumar et al., 2012). Yet, not all microbes have the same effects. For example, while some microbes bioadsorb and trap heavy metals, thus preventing them from entering plant cells (Madhaiyan et al., 2007), others produce siderophores, which often increase uptake of heavy metals by plants (Braud et al., 2009). There is great potential to harness this variation in microbial effects by optimizing plant-microbe associations for bioremediation of contaminated sites (Glick, 2003; Rajkumar et al., 2012).

The extreme evolvability of microbial communities may explain why they underlie so much variation in plants. High species diversity, large population sizes, short generation times, and horizontal gene transfer may increase the response to selection in microbes relative to their plant hosts (Polz et al., 2013; Mueller and Sachs, 2015). For example, articial selection on root-associated microbes resulted in 50% later flowering in *Arabidopsis thaliana* (Panke-Buisse et al., 2015). Furthermore, in dual-selection on plants and soil biota for plant drought tolerance, it was microbial species turnover, and not plant adaptation, that accounted for most of the increase in drought tolerance (Lau and Lennon, 2012). Dual-selection on plants and microbes could involve microbe-independent responses in plants, plant-independent responses in microbes, or co-dependent responses, such as local adaptation between plants and microbes for the ability to receive more benefit from local partners or favor beneficial partners (Kiers et al., 2003; Johnson et al., 2010; Lundberg et al., 2012; Simonsen and Stinchcombe, 2014; Wagner et al., 2014; Batstone et al., 2016; Rúa et al., 2016). However, when microbiomes are not perfectly transmitted from parent to offspring, micriobial fitness is not perfectly aligned to plant fitness or selection by breeders, and individual microbes may evolve to their own benefit at the expense of hosts (Douglas and Werren, 2016). Indeed, individual microbes can be responsible for dramatic increases or decreases in plant fitness (Berg and Smalla, 2009).

Duckweed has been of long and continuing interest for its ability to take up a wide variety of contaminants from water (Mo et al., 1989; Stout et al., 2010; Stout and Nüsslein, 2010; Sekomo et al., 2012; Gatidou et al., 2017). Duckweed have potential for bioremediation of many contaminants, including low-level zinc contamination (Dirilgen and Inel, 1994; Radić et al., 2010; Jayasri and Suthindhiran, 2017), a common problem of urban and suburban water bodies (Göbel et al., 2007), including around Ontario (Miller et al., 1992; Glooschenko et al., 1992; Liskco and Struger, 1996; Ontario Ministry of the Environment, 2011), likely having moderate detrimental effects on animal life (Glooschenko et al., 1992; Miller et al., 1992; Heijerick et al., 2002). Duckweed lines are known to vary in sensitivity to zinc concentrations (Van Steveninck et al., 1990), though tolerance may depend also on the presence of other heavy metals (Dirilgen and Inel, 1994; Balen et al., 2011). Duckweed growth rate is the primary factor influencing contaminant removal (Zhao et al., 2014a), and this varies across duckweed genotypes and nutrient levels (Sree et al., 2015; Ziegler et al., 2015).

It is unclear how much of our understanding of plant-microbe interactions in terrestrial environments translates to floating aquatic habitats. For example, unlike the rhizosphere microbiomes of terrestrial plants, the diversity of species growing with *Lemna minor* is likely much lower (Ishizawa et al., 2017b). However, there is already some suggestive evidence for GxGxE effects in duckweed-microbiome-contaminant systems, in that microbial communities affect duckweed growth rates and contaminant removal (Zhao et al., 2014b, 2015; Ishizawa et al., 2017b). Here, we explored the interactive effects of microbial communities, zinc contamination, and host genotype on duckweed fitness and phenotypes, as well as on microbial growth. We hypothesized that as in terrestrial plants, fitness and phenotypes would be shaped by the interactive effects of biotic and abiotic environments.

## Methods

To investigate the interactive effects of microbial communities and contaminants on duck-weed growth and phenotypes, we tested four populations of duckweed with three microbial treatments in two zinc environments. We collected duckweed from ponds in the Greater Toronto Area in the summer of 2017 (Table 1). We first extracted microbes by pulverizing fresh tissue from each site of 1 or 2 fronds, plating the slurry onto yeast mannitol media agar plates, culturing at 29°C for 5 days, and placing at 4°C for storage. These microbes comprise the subset of the duckweed microbiome that can live on yeast mannitol media. We transferred field-collected live duckweed to growth media (Krazčič et al., 1995) and maintained stock populations of each duckweed line in 500 mL glass jars in a growth chamber with a cycle of 23°C and 150 μmol/m^2^ lighting for 16 hours followed by 18°C and dark for 8 hours.

**Table 1:** Locations of source sites for duckweed and associated microbes.

| Site | Latitude | Longitude |
| --- | --- | --- |
| Kelso | 43.51 | -79.95 |
| McCraney | 43.44 | -79.74 |
| Moccasin | 43.73 | -79.33 |
| Stoney Creek | 44.28 | -78.70 |

Our three microbial treatments were: none, added “home” microbes, and added “away” microbes from one of the other three populations. Which “away” microbes a duckweed population received was selected randomly without replacement. The randomization paired plants from Kelso with microbes from Moccasin, plants from McCraney with microbes from Kelso, plants from Moccasin with microbes from Stoney Creek, and plants from Stoney Creek with microbes from McCraney. We made a separate microbial inocula from each site by taking a swab across the stored agar plate, adding to liquid yeast mannitol media, culturing in a shaker for five days at 30°C and 200 rpm, and then diluting to control optical density across inocula. Because communities are made of various species, the translation of optical density to cell count is inexact (see below), but inocula were approximately 200 cells per μL.

We placed each experimental plant into 2.5 mL autoclaved growth media (Krazčič et al., 1995) in a 3.4 mL well of a 24-well plate. Before adding each plant, we removed surface microbes from the lab culture by dipping in 75% ethanol. Plants selected were approximately the same size, and all had one mature and one immature frond each. After adding plants to wells, 20 μL of the treatment-specific microbial inoculum was added to each well.

We crossed the 12 combinations of duckweed genotype and microbial community with low (0.86 micromolar) or high (3.44 micromolar) zinc water concentrations; four times the amount of ZnSO_4_ was added to the high compared to the low zinc treatment solution (a neglible increase of sulfate concentration). These concentrations of zinc represent natural (Liskco and Struger, 1996) and elevated levels, such as those generated by waste discharge (Ontario Ministry of the Environment, 2011) runoff events (Liskco and Struger, 1996; Göbel et al., 2007). We repeated the full design over 10 replicates per treatment combination for a total of 240 plants in 10 24-well plates. Replicates were assigned to wells at random, and as a result treatments were spatially interspersed within plates.

We photographed plates under a stationary camera at the beginning and end of the experiment. After taking the beginning photograph, each plate was sealed with a gas-permeable membrane to prevent contamination among wells, and placed into a growth chamber set to the same conditions as above (23°C and 150 μmol/m^2^ lighting for 16 hours, 18°C and dark for 8 hours). After 10 days, we removed plates and photographed again to measure change in growth and phenotypes. From the photographs, we hand counted the number of final fronds in each well, and used ImageJ (Schneider et al., 2012) to measure the total pixel area of duckweed from start to finish (growth rate), greenness of fronds (relative to blue and red), and the ratio of the total pixel area of the fronds in a well to the total perimeter of fronds in a well as a measure of the tendency of fronds to aggregate. More aggregated fronds may be more stable and dense on the water surface, which could increase shading of the water and reduce warming. Much local duckweed habitat in the sampling area is stormwater ponds, and reducing temperatures of the outflow water to streams is of concern for fish (e.g. Chu et al., 2005; Herb et al., 2009; Comte et al., 2013). Greenness should be a coarse indication of chlorophyll content (Adamsen et al., 1999; Keenan et al., 2014), indicating potential, rather than realized, growth.

As a measure of microbial growth, we measured the optical density of suspended well solution, using starting growth media in the well as a blank control. Plates were frozen and stored at −20°C before optical density measures, so that microbial growth did not continue as measurements were taken. For a subsample of replicates (3 out of 10), we also measured microbial growth with an alternate method. Immediately after the end of the experiment (before freezing), we sampled 10 μL of well solution, diluted to 1 mL and then plated 10 μL of the dilution onto agar petri dishes and grew at 29°C for 5 days. We then scored the number of colony forming units (CFUs) on plates, as a measure of the total microbial growth. Some plates had colonies too numerous to accurately count, so we excluded those measurements (4 total) from analysis.

We analyzed data in R (R Core Team, 2014). We first quantified the effect of inoculation with microbes on duckweed growth and microbial optical density to verify that manipulation of microbes was successful and that it positively affects duckweed. We used linear models in MCMCglmm (Hadfield, 2010) with change in duckweed pixel area, final frond number (all wells started with 2 fronds), and optical density in wells as the separate response variables (10,000 iterations, 1,000 burn in, thinning by 50). We subset to data for inoculated treatments for the remaining analyses.

For each response variable, we searched for the best model out of all possible models including the potential explanatory variables of duckweed population, microbe population, combination of duckweed and microbes categorized as “home” or “away”, zinc treatment, and all possible two-way interactions using the dredge (package MuMIn Bartoń, 2013) and MCMCglmm functions (10,000 iterations, 1,000 burn in, thinning by 50), limited to a maximum of 4 parameters in addition to the intercept. Best models were determined by comparing DIC (Spiegelhalter et al., 2002). We re-fit the top three models for each response variable with increased MCMC parameters (100,000 iterations, 10,000 burn in, thinning by 500) to verify the best model. If the top 3 models were indistinguishable in DIC (swapped order in DIC fit across repeated MCMC chains), we selected the simplest. Frond number was treated as poisson distributed, all other variables were treated as gaussian. We report results for the best model for each response variable. We determined significant differences between treatments using 95% highest posterior density intervals as calculated from the posterior distribution of parameter effects.

Finally, we asked whether associations exist between duckweed growth measures (plant fitness), traits, and optical density (total microbial growth, putatively linked to average microbial fitness across species) using treatment means. Strong associations would suggest that duckweed fitness is linked to duckweed traits or total microbial abundance. We also verified a relationship between optical density and colony forming units (i.e. live cells) in the subset of wells with both measurements. While both attempt to measure microbial growth during the experiment, colony forming units suffers from post-experiment competitive dynamics on petri plates, overgrowth obscuring some individual colonies, and being laborious data to collect. Optical density is not affected by post-experiment dynamics nor human counting error, but is unlikely to be equal across all microbe species, and may be influenced by dead cells. We measured each association by again fitting linear models with MCMCglmm() using duckweed growth, traits, and optical density as paired response and explanatory variables, and using optical density and CFUs (100,000 iterations, 10,000 burn in, thinning by 500).

## Results

We aimed to understand GxGxE effects on duckweed traits and fitness across duckweed gentoype, microbial communities and contaminant environments. Using a growth chamber experiment, we manipulated all three sources of variation, and quantified the effects of distinct microbial communities and zinc levels across different duckweed populations. We found first that adding microbes had a significant positive effect on average growth measured in either pixels or frond number (both pMCMC < 0.001) relative to sterilized duckweed alone, suggesting that the microbial communities as a whole, or some subset of the species, provide benefits to duckweed in the laboratory environment (Figure 1, across all other treatments combined). We also found that while our sterilization procedure for duckweed fronds was imperfect, there were many fewer microbes detected in the wells where microbes had not been re-inoculated onto duckweed (pMCMC < 0.001, Figure 1).

**Figure 1:**
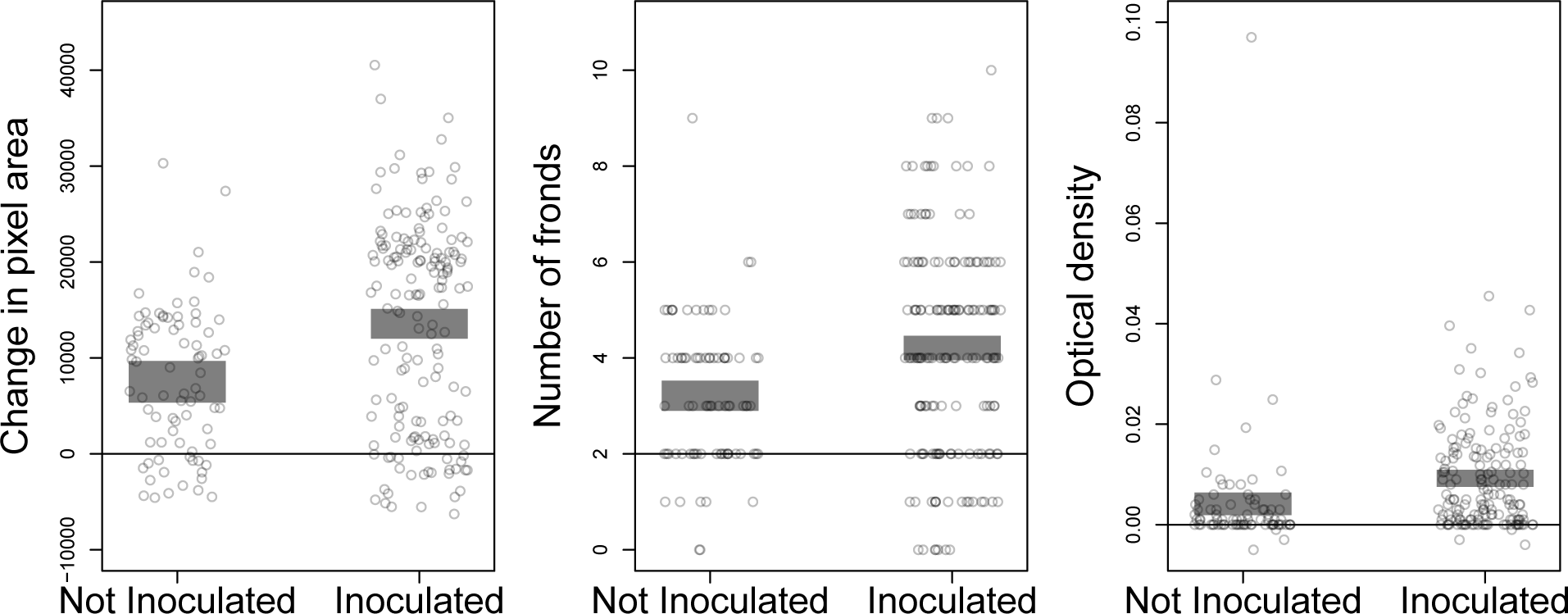
Inoculation with microbes increased duckweed fitness and total microbial growth, compared to surface-sterilized plants that did not receive additional microbes. From left to right, plots show change in pixel area, final duckweed frond number (horizontal line at 2, indicates starting frond number), and optical density of suspended microbes comparing inoculated and uninoculated treatments. Open points are individual wells. Grey regions are 95% highest posterior density intervals for the mean.

We next explored variation in duckweed fitness across experimental treatments. Our best model for growth in pixel area included both variable effects of duckweed source population (pMCMC < 0.01), microbe source population (pMCMC < 0.01) and consistently negative effects of increasing zinc level (pMCMC < 0.001). Comparing 95% highest posterior density intervals (HPDI), duckweed plants from Moccasin and Stoney Creek grew larger than plants from Kelso and McCraney, and plants from McCraney grew less than those from Kelso. Additionally, duckweed plants growing with Moccasin microbes grew more than plants with other microbes, and duckweed with Kelso microbes grew less (95% HPDI, Figure 2, top panel). While the best model did not include interactive effects between any of these terms on growth in pixel area in the best model (i.e. there were no GxG, GxE, or GxGxE effects), we note that duckweed and microbe main effects are dif cult to distinguish from duckweed and microbe interactive effects in this experimental design. The best model for frond number (Figure 2, middle panel) included only effects of duckweed source population (pMCMC < 0.01) and microbe source population (pMCMC *<* 0.01), with nearly identical plant and microbe treatment effects as in the pixel area model, except that plants in Moccasin microbes grew the same number of fronds as plants in microbes from Stoney Creek and McCraney. Biotic context (microbial community), environment (zinc), and plant genotype (source) thus independently influence at least one aspect of duckweed fitness.

**Figure 2:**
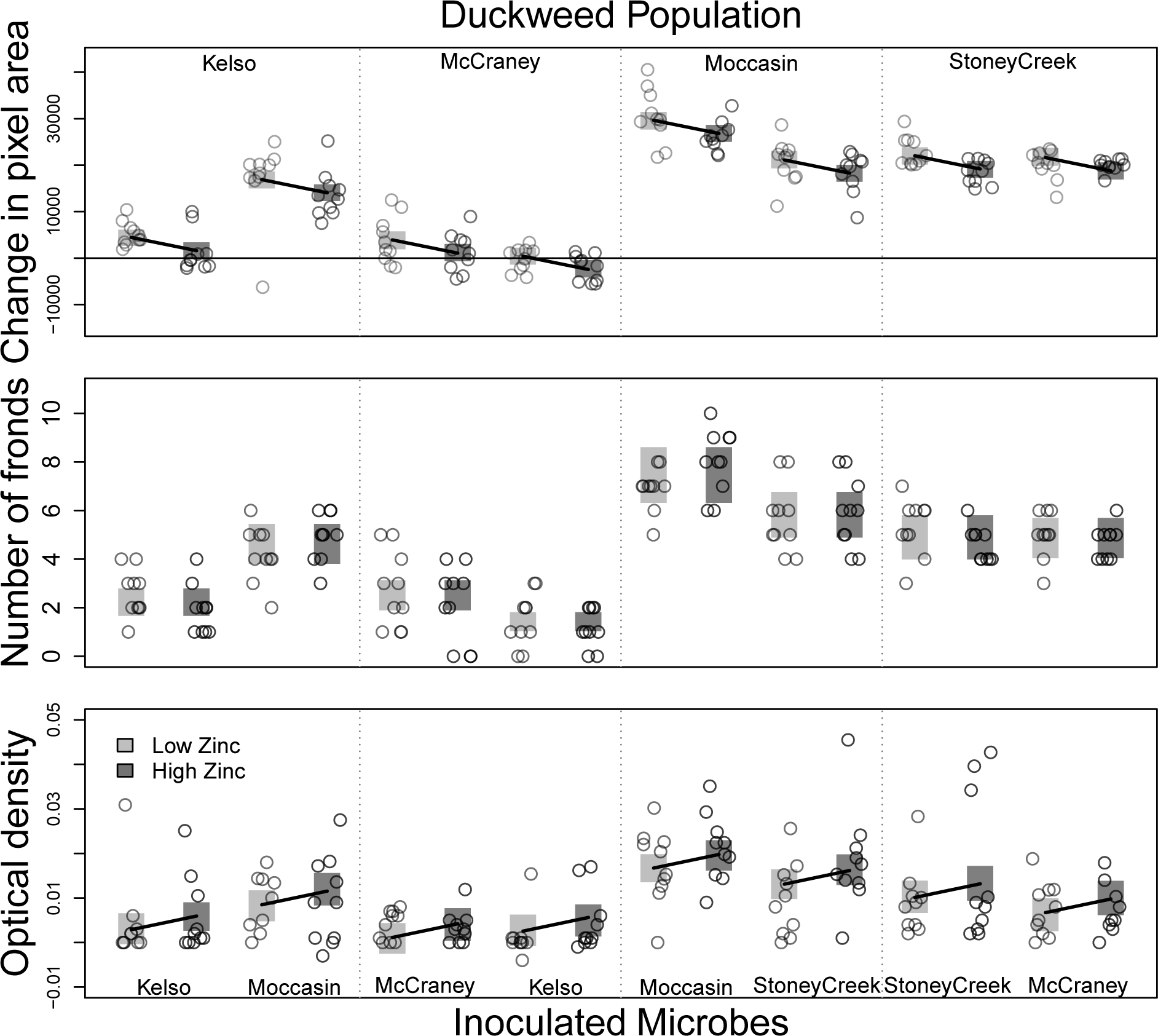
Duckweed fitness and total microbial growth (optical density). Change in pixel area (top panel) and frond number (middle panel) of duckweed plants. Bottom panel shows optical density relative to blank media from samples of each experimental well at the end of the experiment. Points are individual wells. Grey regions are best model 95% highest posterior density intervals for treatment means. Different populations of duckweed are separated by vertical dashed lines and labeled at the top, while microbe treatments are labeled at the bottom. High zinc levels are indicated with darker gray. Plant population and microbe effects are significant in all panels, whereas zinc effects (indicated by solid black lines), are significant for pixel area change and optical density only.

Total microbial growth, inferred from optical density, also varied across experimental treatments. The best model for optical density included effects of microbe source (pMCMC < 0.01), plant source (pMCMC < 0.01), and zinc treatment (pMCMC < 0.05). Wells inoculated with microbes from Moccasin had the highest microbial growth, and plants from Stoney Creek and Moccasin supported greater increases in optical density, as did treatment with high zinc (all at 95% HPDI, Figure 2). These results suggest that microbial growth is altered by variation among plant inputs, microbial community composition, and zinc runoff levels, but not altered by interactions among these biotic and abiotic influences. We also explored the relationship between optical density and colony forming units (i.e. viable cells) in the subset of wells for which we have both measures. The two measures are moderately correlated with marginal significance (*ρ* = 0.28, pMCMC < 0.1, Figure S1).

To assess whether treatments have concerted or separate effects on microbial growth and plant fitness, we fit models of treatment mean fitness or growth for plants and microbes. Duckweed fitness was significantly related to microbial growth (effectively, fitness averaged across all species in optical density measures) for both pixel area (pMCMC < 0.01) and frond number (pMCMC < 0.001, Figure 3). This suggests strong fitness alignment, with microbes that provide duckweed with fewer fitness benefits also having lower fitness, and vice versa. This occurs despite positive responses to zinc among microbes, but negative responses in duckweed (see above, Figure 2). However, fitness alignment should be interpreted with caution because abundance was not measured separately for each microbe species.

**Figure 3:**
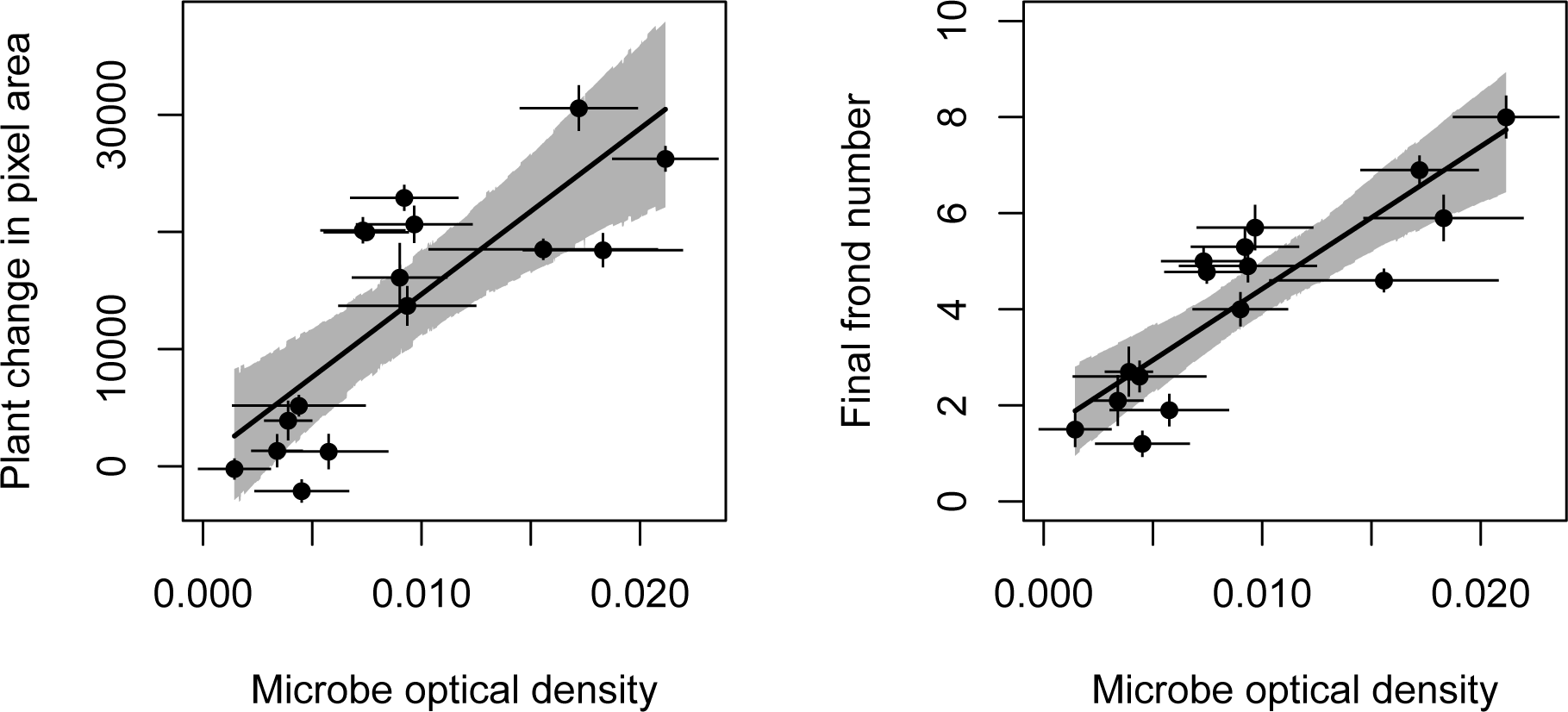
Fitness alignment between duckweed and microbes across experimental treatments, where duckweed fitness is measured either as increase in pixel area or final frond number and microbe “fittness” is optical density measured across all species in the community. Points are means for each experimental treatment, and whiskers are standard errors. The linear relationships in the background are the model predictions for the means (solid line) with 95% highest posterior density intervals in gray.

The factors best explaining variation in phenotypes differed between plant phenotypes. For greenness of floating tissue, plant population was the only explanatory variable included in the best model (pMCMC < 0.01), and plants from Moccasin and Stoney Creek were greener than others (95% HPDI, Figure 4). In contrast, the best model for frond aggregation (pixel area divided by the perimeter of all particles) included microbe (pMCMC *<* 0.01) and plant source effects (pMCMC < 0.01), as well as negative effects of increased zinc (pMCMC < 0.05). Duckweed from McCraney were significantly less aggregated than other duckweed, while duckweed from Moccasin and Stoney Creek were more aggregated, and microbes from Kelso supported less aggregated duckweed plants (95% HPDI, Figure 4). In sum, we see that plant genotype (source), biotic interactions (microbial community), and aquatic toxins (zinc level) affect phenotype expression unequally across phenotypes.

**Figure 4:**
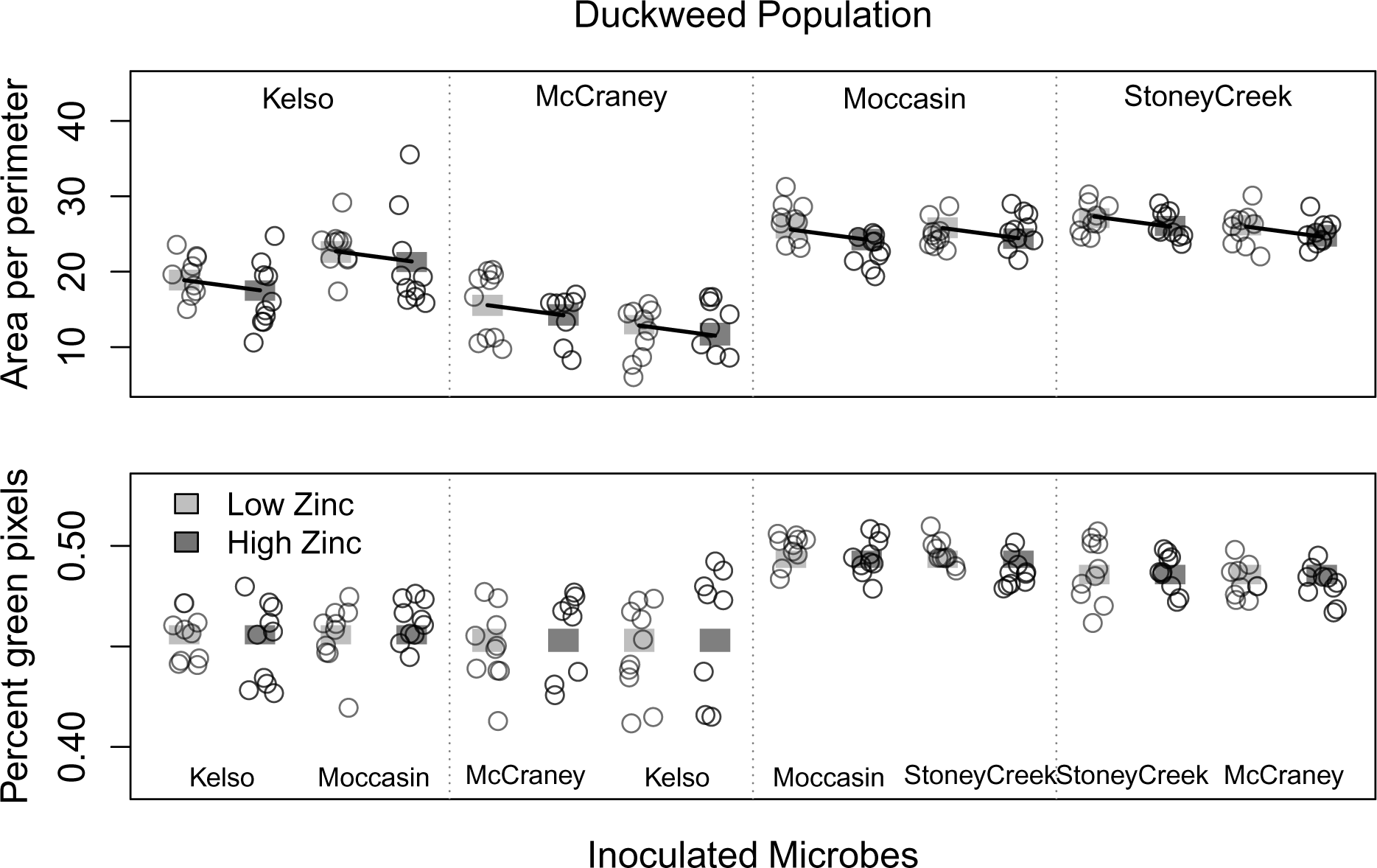
Duckweed traits. Frond aggregation (top, ratio of total pixels to total perimeter) and greenness (bottom, percent of pixel area that is green) in duckweed plants at the end of the experiment. Points are individual wells. Vertical blocks behind points are best model predicted 95% highest posterior density intervals for the means of each treatment combination. Different populations of duckweed are separated by vertical dashed lines and labeled at the top, while microbe treatments are labeled at the bottom. High zinc levels are indicated with darker gray. Plant population explains significant variation for both traits, whereas aggregation is additionally explained by microbe and zinc level (model zinc effect depicted by solid black lines).

Finally, we observed some co-correlations among response variables. Growth in pixel area is correlated with frond number (*ρ* = 0.88), frond aggregation score (*ρ* = 0.84) and greenness (*ρ* = 0.69). Likewise, greenness is correlated with frond number (*ρ* = 0.72) and aggregation score (*ρ* = 0.70), and aggregation score is also correlated to frond number (*ρ* = 0.70). Microbial abundance (optical density) is also correlated with greenness (*ρ* = 0.40), and aggregation score (*ρ* = 0.36), indicating a link between duckweed phenotypes, duckweed fitness, and microbe growth (all slopes significant at pMCMC < 0.01).

## Discussion

Microbial communities living near, on, and inside host tissues constitute a ubiquitous aspect of the biotic environments of plants. Just like abiotic conditions, microbial biotic conditions can affect the expression of phenotypically plastic traits and fitness in terrestrial plants (Friesen et al., 2011; Wagner et al., 2014). We explored how abiotic and biotic factors may together or separately influence trait development and fitness in duckweed and its associated microbes. We found strong differences in phenotypes and fitness across duckweed driven by duckweed origin, the origin of co-cultured microbial communities, and treatment with the aquatic contaminant zinc, but no co-dependent effects.

Interestingly, while the effect of duckweed source population affected all phenotypes and fitness measures, and microbe source affected most measures, only for pixel area, optical density, and aggregation did we observe effects of abiotic environments (Figures 2,4). The contrasts among patterns for phenotypes and fitness of duckweed is somewhat surprising, since growth in area and number of individuals should both be measures of growth rate, and since both measured phenotypes are presumably linked to fitness (all have significant pairwise correlations). Greenness should be primarily related to chlorophyll content and future reproductive potential, and aggregation is likely the inverse of vulnerability to air and water current dispersion. It could be that the lab environment prevents fitness effects of variation in these phenotypes, or that other, unmeasured, phenotypes dominate effects on fitness. Alternatively, we may have limited power to quantify abiotic and interactive effects on fitness due to dramatic main effects of duckweed and microbial sources, incomplete culture of the field microbiome under lab conditions, or incomplete sterility before microbial inoculation, although strong effects of microbial inoculation (Figures 1,2,4) suggests minimal influence of incomplete sterility.

Genetic diversity among duckweed populations is a possible source for the significant variation across duckweed population sources. However, existing work suggests fairly low genetic diversity in *L. minor* in the local region (Ho, 2017). Duckweed phenotypic diversity could also come from variation in endosymbiotic microbes, which would not have been removed by our surface sterilization, or from epigenetic differences across populations. Such genetic, epigenetic, or endosymbiotic diversity might be generated by neutral divergence among populations, or by trade-offs for phenotypes across environments (e.g. Prati and Schmid, 2000; Agrawal et al., 2010; Albert et al., 2010), both commonly observed phenomena.

The substantial phenotypic and growth differences among microbial treatments that we observed are likely in large part due to differing microbial species composition, because the effects of microbial communities on plants are often highly contingent on community composition (Berg and Smalla, 2009). Interestingly, community composition itself can be a function of plant influences (e.g. microbially driven plant-soil feedback Klironomos, 2002; Anacker et al., 2014). The underlying question for the effects we observed here is thus why microbial communities may differ across sites. Environmental filtering of species, random colonization differences across space, and the duckweed plants themselves (e.g. plant-water feedbacks) could be involved in generating these different communities. Analogous to microbially driven plant-soil feedbacks observed in terrestrial plants (Bailey and Schweitzer eds., 2016), such plant-water feedbacks could be common. Consistent with potential for plant-water feedbacks, duckweed sources seem to drive the overall increases in microbial abundance (Figure 2), however, it remains unknown whether duckweed plants influence microbial community composition.

The microbial communities investigated here can best be described as beneficial (Figure 1) from the perspective of the duckweed. In plant-microbe mutualisms, we generally see positive correlations between host and symbiont fitness (Friesen, 2012), although some environments may decouple them (Weese et al., 2015; Shantz et al., 2016). Aquatic microbes associated with duckweed species that may affect growth are known to include diatoms (Desianti, 2012), nitrogen-fixing cyanobacteria (Zuberer, 1982; Duong and Tiedje, 1985; Eckardt and Biesboer, 1988), and a collection of additional bacteria, including members of other nitrogen-fixing clades (Underwood and Baker, 1991; Ishizawa et al., 2017b), and one that may provision phosphorus (Ishizawa et al., 2017a). Here we find positive correlations between duckweed fitness and microbial growth across treatments (Figure 3), potentially indicating positive fitness feedbacks (Sachs et al., 2004) between duckweed and the community of microbes that live on them. This positive fitness association is despite average decreases in duckweed fitness, and average increases microbial growth, in response to increased zinc, and suggests that zinc in runoff water will not cause mutualism breakdown between duckweed and microbes.

The differences across duckweed populations and microbial communities we see here may alter the potential of duckweed to remediate environments contaminated with zinc or other pollutants. Others have postulated that duckweed may be of interesting and unique value in phytoremediation of water (Mkandawire and Dudel, 2007; Ziegler et al., 2016), specifically due to its uptake or modification of a wide variety of aquatic pollutants (Mo et al., 1989; Stout and Nüsslein, 2010; Stout et al., 2010; Sekomo et al., 2012; Uysal, 2013; Sasmaz et al., 2015; Baciak et al., 2016; Gatidou et al., 2017; Gomes et al., 2017). Plant-associated microbes are often in part responsible for removal or detoxification of contaminants, and presence of various taxa on duckweed may alter its phytoremediation potential (Toyama et al., 2009; Zhao et al., 2015). Microbes may alter phytoremediation through impacts on plant growth rate (Glick, 2003; Sobariu et al., 2017), by altering the relative rates at which non-toxic nutrients and toxic pollutants are taken up (Burd et al., 2000), or by directly metabolizing or altering pollutants, as has been discovered in a microbe inhabiting the roots of another duckweed species (Toyama et al., 2009). Here, we focused on zinc contamination. Zinc was previously found to both be sequestered by duckweed, and to physiologically affect duckweed (Radić et al., 2010; Jayasri and Suthindhiran, 2017). We found that microbes from different natural duckweed sites alter duckweed growth rates, respond positively to zinc, and generally increase duckweed fitness (Figures 1,2, and 3). Thus microbes likely indirectly influence the ongoing and potential amount of phytoremediation in duckweed-inhabited sites. However, how microbiomes affect the fate of zinc or other contaminants, and whether microbiome species composition plays a predictable role remain open questions.

## Conclusions

Here, we found that microbiome variation has complex effects on phenotypes and fitness in an aquatic plant, similar to how microbiome variation affects terrestrial plants. This is despite the fact that duckweed draws a microbiome from the water environment that is less complex than typical terrestrial plant microbiomes (Lundberg et al., 2012; Ishizawa et al., 2017b). As a smaller plant with a simpler microbiome, more manipulative experimentation is possible for duckweed microbiomes than for terrestrial plant microbiomes. We expect that duckweed and its associated microbiome will thus prove pivotal in experimentally elucidating properties of ecology and evolution in plant-microbiome interactions, and in manipulating these effects for applied approaches, such as phytoremediation.

## Acknowledgements

This work was funded by the Natural Sciences and Engineering Council of Canada (NSERC), through a Discovery Grant to MEF (RGPIN-2015-06742) and a Canada Graduate Scholarship to JL. EL was supported by the University of Toronto Centre for Global Change Science. The authors would like to thank D. Sinton and B. Nguyen for engineering solutions improving our experimental set-up and members of the Frederickson lab for discussion. JL, EL, and MEF executed collections. AMO and JL ran the experiment and collected data. AMO performed analyses and provided the first draft. All contributed to study design, revised the manuscript, and gave approval for publication.

---

**Figure S1:**
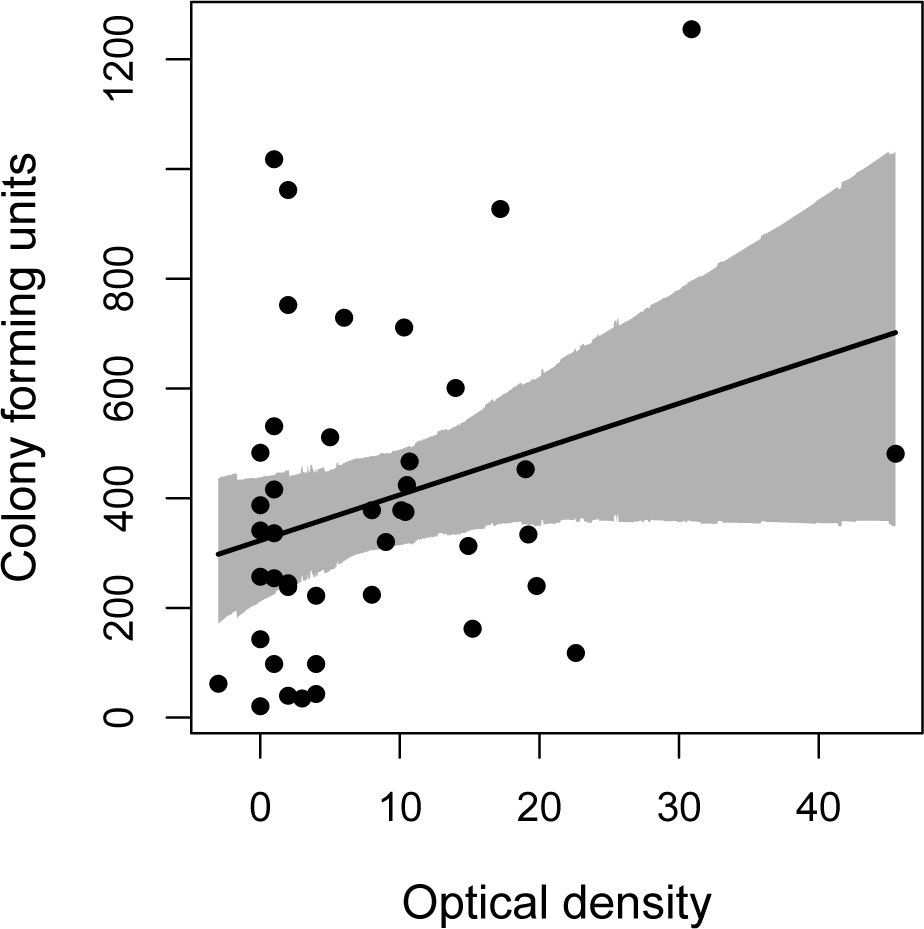
Correlation between microbial community fitness measures. Points are a subset of experimental wells for which both colony forming units and optical density were measured. The linear relationship in the background is the model predictions for the mean with 95% highest poserior density intervals.

